# Minimal determinants for lifelong antiviral antibody responses in BALB/c mice from a single exposure to virus-like immunogens at low doses

**DOI:** 10.1101/2023.02.20.529089

**Authors:** Wei-Yun Wholey, Alexander R. Meyer, Sekou-Tidiane Yoda, Bryce Chackerian, Julie Zikherman, Wei Cheng

**Author notes:** Corresponding author: Dr Wei Cheng University of Michigan 428 Church Street Ann Arbor, MI, 48109-1065. Both authors contributed equally to this work (Wei-Yun Wholey, Alexander R. Meyer).

## Abstract

The durability of an antibody (Ab) response is highly important for antiviral vaccines. However, due to the complex compositions of natural virions, the molecular determinants of Ab durability from viral infection or inactivated viral vaccines have been incompletely understood. Here we used a reductionist system of liposome-based virus-like structures to examine the durability of Abs in primary immune responses in mice. This system allowed us to independently vary fundamental viral attributes and to do so without additional adjuvants to model natural viruses. We show that a single injection of antigens (Ags) orderly displayed on a virion-sized liposome is sufficient to induce a long-lived neutralizing Ab (nAb) response. Introduction of internal nucleic acids dramatically modulates the magnitude of long-term Ab responses without alteration of the long-term kinetic trends. These Abs are characterized by exceptionally slow off-rates of ∼0.0005 s^-1^, which emerged as early as day 5 after injection and these off-rates are comparable to that of affinity-matured monoclonal Abs. A single injection of these structures at doses as low as 100 ng led to lifelong nAb production in BALB/c mice. Thus, a minimal virus-like immunogen can give rise to potent and long-lasting antiviral Abs in a primary response in mice without live infection. This has important implications for understanding both live viral infection and for optimized vaccine design.

## Introduction

The immune system robustly mounts effective Ab responses against a vast range of viral pathogens. This property of the immune response forms the biological basis for many of the vaccines that have been licensed for use in the US. Despite the recent great success of COVID-19 vaccines, there remain challenges that face vaccine development. Among these, a major issue is the durability of protection. At a mechanistic level, conditions that promote the durability of nAb responses are not well understood^1^, and the rules that govern the durability of vaccine protection are yet to be defined.^1^ Some vaccines, such as those against measles, mumps, and rubella, elicit Abs whose plasma titer half-lives span more than 100 years^2^; in contrast, nAb response wanes quickly even after a second dose of the SARS-CoV-2 mRNA vaccine^3-5^, which highlights the importance of booster vaccines for continued protection.

Why do mRNA vaccines require continued boosters to achieve durable protection while certain live attenuated vaccines such as measles and mumps can lead to lifelong protection^2^ after only two injections? Currently, we don’t have complete answers to this question. However, it has been long recognized that the biophysical form of an Ag is likely to be a critical factor governing Ab durability^6-8^. It has been well documented that multivalent virus-like particles (VLPs) or whole-virus vaccines can induce durable and protective IgG in both preclinical^9^ and clinical settings^7^. Prior studies also established that Ag valency has profound impacts on Ab responses compared to monovalent Ags^10-12^, including the ability of B cells to differentiate into effector cells^11^, clonotype diversity and neutralization breadth^12^. Recently, we have also shown – by contrast to monovalent soluble Ag – that multivalent Ag display on virus-sized liposomes alone, even at a low epitope density, can trigger robust Ag-specific B cell signaling, survival, and proliferation by evading Lyn-dependent inhibitory pathways for B cell activation^13^. These results indicate that much remains to be learned about mechanisms whereby the biophysical form of an Ag influences the durability of an Ab response.

To understand mechanisms of Ab durability brought by multivalent Ags such as VLPs or whole viruses, one major barrier in our view is the complex ingredients of commonly studied VLPs, as well as the inherent complexity in studies of viral infections, where inflammation^14,15^ and cytokine secretion^16^ can modulate the immune network to confound Ab responses. Also, for almost all vaccine studies in literature, different adjuvants have been extensively used to induce effective immunity. Under these circumstances, it is difficult to unravel the mechanisms of Ab durability induced by multivalent Ags because the mechanisms of adjuvants have not been completely defined in many cases.

To overcome these limitations, we have developed a reductionist system of synthetic virus-like structures (SVLS) based on liposomes to mimic the complex biophysical features of enveloped viruses^17, 18^. These SVLS are constructed using highly purified biochemical ingredients so that the molecular basis of their immunogenicity is well defined. These SVLS are highly modular in their supramolecular structures so that we can dissect the contributions of critical viral features to the durability of Ab responses by modulating each one independently and collectively. Here, we show that multivalent display of a protein Ag alone on virion-sized liposomes can induce long-lasting production of IgG in mice upon a single immunization. Inclusion of internal nucleic acids (iNA) in these structures profoundly modulates the magnitude of IgG in both the immediate and long-term so that a single injection of SVLS at a very low Ag dose can lead to lifelong Ab production in BALB/cJ mice. The Ag-specific IgGs from SVLS immunization are characterized by ultralow off-rates and offer potent and long-lasting neutralization activity. Therefore, these results reveal minimal molecular determinants for a durable Ab response previously observed from live viral infections. The enduring Ab responses as we observed in two entirely different and unrelated protein Ags indicate that our findings may have general relevance for understanding viruses and developing vaccines.

## Results

### Four sets of SVLS chosen for current immunization studies

To understand the minimal determinants for durable B cell antibody responses towards virus-like immunogens, we have recently engineered and characterized several sets of SVLS for the current study^17-19^. These SVLS were assembled completely using highly purified proteins, lipids and nucleic acids (NAs), for which the molecular basis of immunogenicity is well defined (Materials and Methods). Two of them, schematically shown on the left in Fig. 1 and termed pRBD (Fig. 1a) and pHEL (Fig. 1c), respectively, are SVLS that display either purified receptor-binding domain (RBD) of the SARS-CoV-2, the causative agent of the COVID-19 pandemic^20^; or a purified mutant version of hen egg lysozyme (HEL), a well-characterized protein Ag that has been used extensively in immunological studies^21^. RBD and HEL are not related to each other in evolution, nor do they possess any significant sequence or structural similarities. Each protein retains their conformations and was displayed in a specific orientation at programmed epitope density on the surface of unilamellar liposomes. To mimic naturally occurring viruses, we intentionally selected lipids that were abundant in natural plasma membranes to construct SVLS (Materials and Methods) and avoided cationic lipids that are known to be immunostimulatory in nature^22^. The interior of pRBD and pHEL was filled with phosphate buffered saline (PBS). The other two sets: pRBD(DNA1) and pHEL(DNA1), schematically shown on the right in Fig. 1, are SVLS that not only display RBD or HEL on their surface, but also encapsulate DNA1, a 20-mer single-stranded DNA that harbors two unmethylated CpG dinucleotides (Materials and Methods). All protein display was achieved through site-specific covalent conjugation on the lipid bilayer surface to ensure stability of the attachment with time^17, 18, 23^. No proteins other than RBD or HEL were present in these structures.

**Figure 1.**
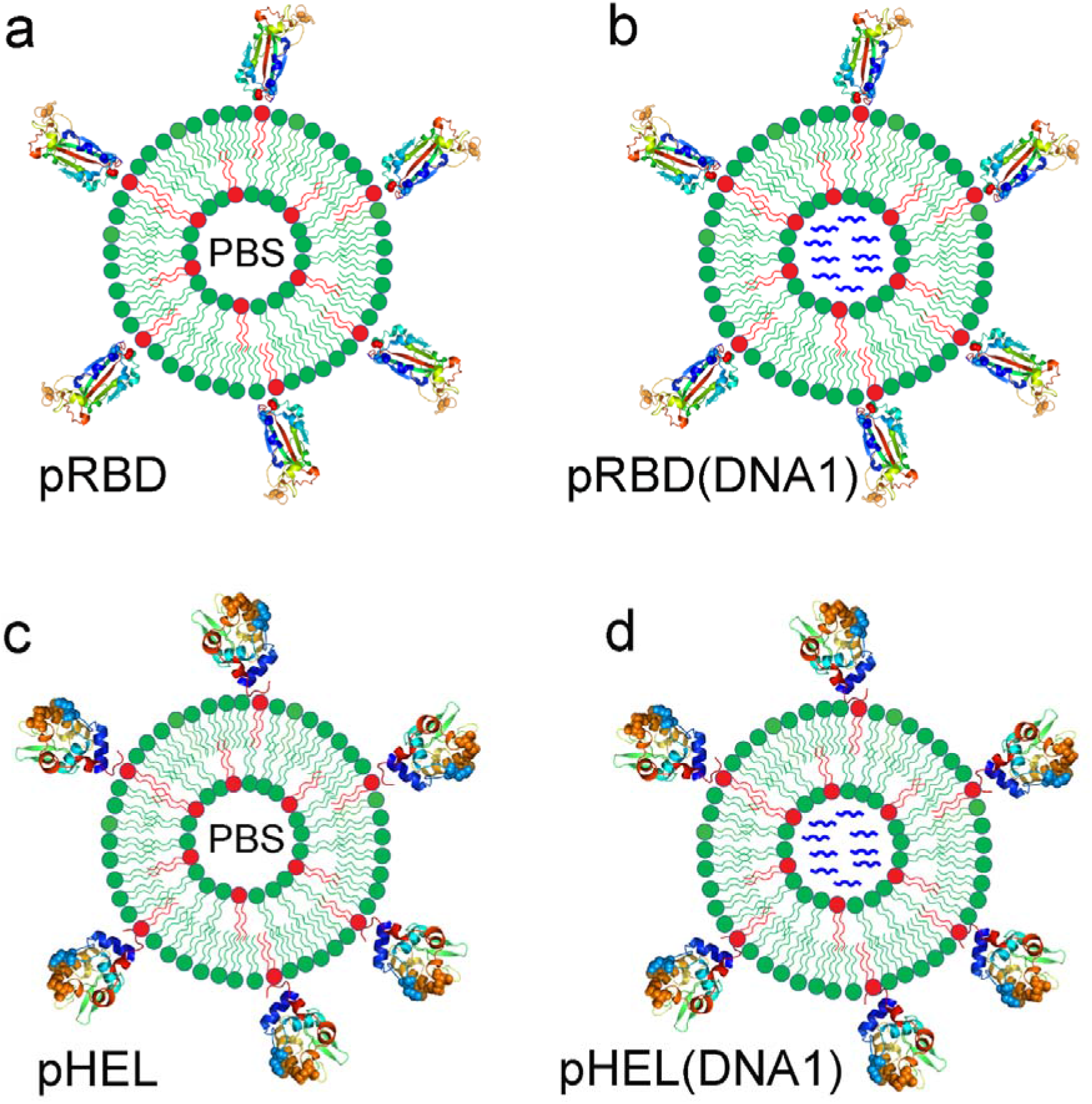
Schematic of four sets of synthetic virus-like structures (SVLS) used in current study that qualitatively and quantitatively incorporate essential features of enveloped viruses. These structures use unilamellar liposomes as the backbones. The surface is conjugated with protein Ags site-specifically via maleimide chemistry at programmed epitope densities. The internal space of the liposome is either PBS as in pRBD (a) and pHEL (c) or loaded with single-stranded DNA oligo DNA1 as in pRBD(DNA1) (b) and pHEL(DNA1) (d) to mimic the NA genomes inside typical biological viruses. The ribbon diagram of RBD used the RBD structure in PDB 6m0j^24^ and the ribbon diagram of HEL used the HEL structure in PDB 1c08^25^.

Most viruses infect their hosts through peripheral routes^26^. To simulate a natural viral infection, we conducted subcutaneous injection of SVLS to deliver a submicrogram dose of Ag into mice. This allows Ag collection by the initial lymphatics to encounter B cells in the draining lymph nodes^27^. A submicrogram dose of Ag contains picomoles of SVLS, which is a very low dose compared to those typically used for VLPs or other conventional model immunogens in mice^9, 28-30^. Importantly, because natural viral infection does not involve any conventional adjuvants such as alum, in current studies, all the injections were conducted without addition of any exogenous adjuvants. Consequently, the injections of SVLS only introduced the highly purified protein Ags and lipids for pRBD or pHEL, and additional encapsulated DNA1 for pRBD(DNA1) or pHEL(DNA1). To focus on primary immune responses, throughout, only a single subcutaneous injection was administered for each mouse.

### SVLS induce durable IgG responses

In the absence of DNA1, we found that a single injection of pRBD at a submicrogram Ag dose is sufficient to elicit long-lived RBD-specific IgG responses in both C57BL/6 (B6) (Fig. 2a triangles) and BALB/c mice (Fig. 2b triangles), although with different kinetic trends. As shown in Fig. 2a, the RBD-specific IgG in B6 mice elicited by pRBD peaked around Day 40 (D40) after the exposure and then decayed over time (triangles). With internal DNA1 (blue symbols), the IgG responses were at least 10-fold higher in magnitude, peaked around two months after exposure, and then slowly decayed. For all cases in Fig. 2a and 2b, the level of RBD-specific IgG in animal sera over 300 days after the single injection of various SVLS remained well above the detection limit of the current ELISA, with the lowest concentration of RBD-specific IgG detected as 36±5 ng/ml for sera collected from B6 mice on D347 after a single immunization with pRBD2 (Fig. 2a, upper triangles). The diamonds in Fig. 2a represent the control sera from B6 mice immunized with an irrelevant immunogen pHEL(DNA1)1. This control demonstrates that the IgG responses are specific towards RBD but not to other components in SVLS such as lipids.

**Fig. 2.**
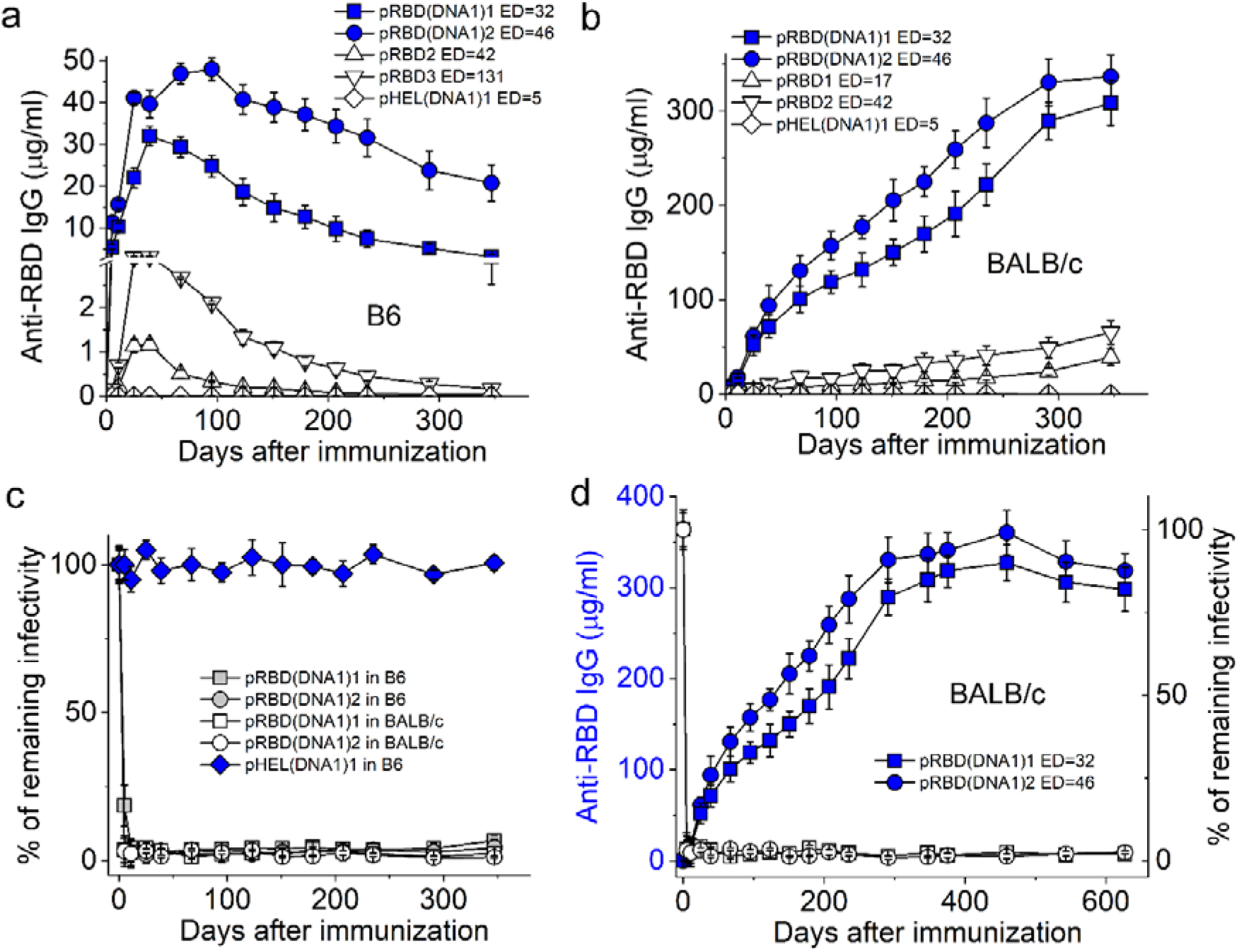
Duration of the IgG response induced by SVLS conjugated with RBD. (**a**) and (**b**) Concentrations of RBD-specific IgG in mouse sera from B6 (a) or BALB/c mice (b) collected over 300 days after a single injection of various agents listed in each inset, with the epitope density for each SVLS listed. All IgG concentrations were measured using ELISA based on standard curves obtained from a reference monoclonal IgG mAb1. (**c**) Time courses of pseudovirion neutralization by various sera from immunized animals as listed in the inset. (**d**) RBD-specific IgG responses induced by pRBD(DNA1) in BALB/c mice over 600 days. Left y-axis and blue solid symbols: concentrations for RBD-specific IgG as measured from ELISA for mouse sera collected on different days after a single injection of the respective agents in BALB/c mice as listed in the inset. Right y-axis and white symbols: time courses of pseudovirion neutralization by mouse sera from BALB/c mice after a single injection of SVLS, with white squares for pRBD(DNA1)1 (ED=32) and white circles for pRBD(DNA1)2 (ED=46). Throughout Fig. 2, the doses of Ags for SVLS displaying RBD were 0.24 µg per animal. N=4 for each time point.

As shown in Fig. 2b, both pRBD (triangles) and pRBD(DNA1) (blue symbols) induced long-lasting IgG whose concentrations in mouse sera increased with time in BALB/c mice, which is in contrast to those in B6. The diamonds in Fig. 2b represent the control sera from BALB/c mice immunized with the irrelevant immunogen pHEL(DNA1)1, demonstrating the specificity of the IgG responses. With internal DNA1, the concentrations of RBD-specific IgG (blue symbols in Fig. 2b) throughout this period were much higher than those induced by pRBD without iNA (triangles in Fig. 2b), like those in B6 and consistent with the notion that innate sensing of NAs regulates adaptive immunity^31, 32^.

To assess the functionality of the RBD-specific IgG measured by ELISA, we prepared HIV-1 based pseudovirions that display the full-length S protein of the SARS-CoV-2 (Materials and Methods). This pseudovirion-based reporter system has been rigorously compared with others and validated as an effective approach for quantitative assessment of serological immunity against SARS-CoV-2^33^. As shown in Fig. 2c, sera collected from these immunized animals over this 1-year span neutralized more than 95% of HIV-1 pseudovirions that displayed the cognate S protein of the SARS-CoV-2 *in vitro*, which included both B6 (grey symbols) and BALB/c mice (white symbols), even though the RBD-specific IgG in B6 waned over this period of time (Fig. 2a, blue symbols). Control sera from B6 immunized with pHEL(DNA1)1 showed no neutralization of virion infectivity (Fig. 2c, blue diamonds), demonstrating Ag specificity of the neutralizations observed. These results suggest that the serum Abs induced by SVLS would be functional *in vivo* to offer the host effective protection against infection of SARS-CoV-2. Continued monitoring of the BALB/c mice immunized with pRBD(DNA1) showed that these nAbs lasted over 600 days (Fig. 2d, blue symbols), approaching the life span of these animals recorded in literature^34^ and the sera remained highly effective in neutralization of cognate pseudovirions throughout (Fig. 2d, white symbols). Despite being different proteins, like RBD, qualitatively similar anti-HEL IgG responses were observed in B6 (Fig. 3a) and BALB/c mice (Fig. 3b) upon a single injection of pHEL or pHEL(DNA1) at a submicrogram Ag dose.

**Fig. 3.**
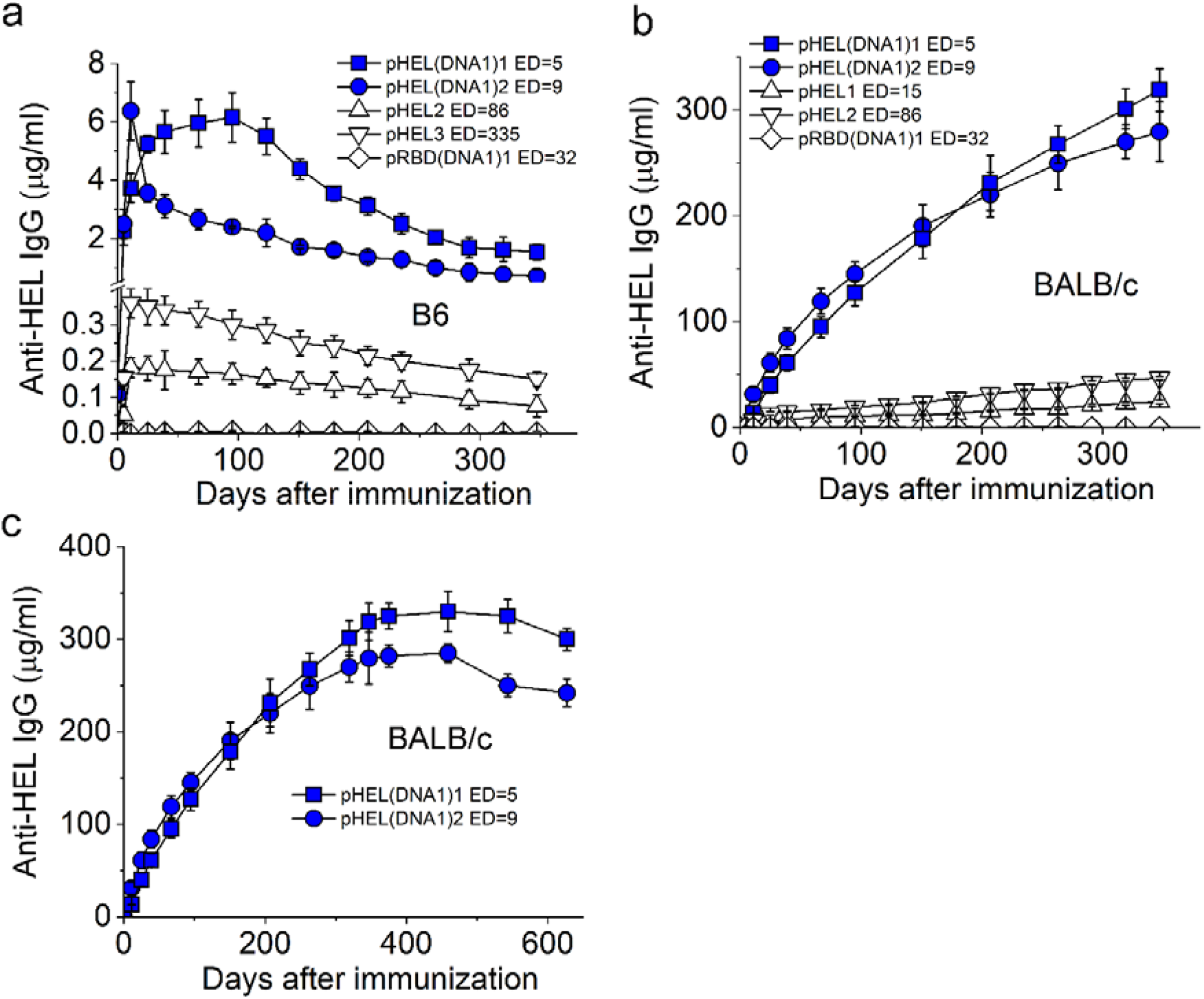
Duration of the IgG response induced by SVLS conjugated with HEL. (**a**) and (**b**) Concentrations of HEL-specific IgG in mouse sera from B6 (a) or BALB/c mice (b) collected over 300 days after a single injection of various agents listed in each inset, with the epitope density for each SVLS listed. (**c**) Concentrations for HEL-specific IgG in BALB/c mouse sera collected over 600 days after a single injection of SVLS as listed in the inset. Throughout Fig. 3, all IgG concentrations were measured using ELISA based on standard curves obtained from a reference monoclonal IgG mAb2. The doses of Ags for SVLS displaying HEL were 0.3 µg per animal for pHEL and 0.1 µg per animal for pHEL(DNA1), respectively. N=4 for each time point.

For all cases in B6 mice (Fig. 3a), the levels of HEL-specific IgG rose initially, which were then followed by slow waning over time. Despite these decays with time, the concentrations of anti-HEL IgG in mouse sera remained well above the limit of detection by the current ELISA. The diamonds in Fig. 3a represent the control sera from B6 mice immunized with an irrelevant immunogen pRBD(DNA1)1, demonstrating that the IgG response is specific towards HEL but not to other components in SVLS such as lipids. It is worth noting that the peak concentrations for HEL-specific IgG induced by pHEL(DNA1) (blue symbols in Fig. 3a) were much lower than those for RBD-specific IgG induced by pRBD(DNA1) (blue symbols in Fig. 2a), likely due to the absence of cognate T cell help for HEL in B6 mice.

As shown in Fig. 3b, both pHEL (triangles) and pHEL(DNA1) (blue symbols) induced long-lasting HEL-specific IgG that increased with time in BALB/c mice, similar to those trends for pRBD and pRBD(DNA1) in BALB/c mice (Fig. 2b) but in contrast to those in B6 (Fig. 3a). The diamonds in Fig. 3b represent the control sera from BALB/c mice immunized with the irrelevant immunogen pRBD(DNA1)1, demonstrating the specificity of the IgG responses. Also, with internal DNA1, the concentrations of HEL-specific IgG (blue symbols in Fig. 3b) were much higher than those induced by pHEL without iNA (triangles in Fig. 3b), analogous to the situation in Fig. 2b for pRBD(DNA1) in comparison to pRBD in BALB/c mice over the same period of time. Like that of pRBD(DNA1), continued monitoring of the BALB/c mice immunized with pHEL(DNA1) showed that these HEL-specific IgG lasted over 600 days (Fig. 3c), suggesting that this lifelong IgG may be a general property that can be achieved for SVLS(DNA1) bearing different foreign Ags in BALB/c mice. For both strains of mice, the presence of internal DNA1 induced a stronger and more durable IgG response compared to SVLS without iNA. In particular, the durable IgG response from pHEL(DNA1) in B6 mice (Fig. 3a blue symbols) suggests that long-lived plasma cells may have been generated in the absence of cognate T cell help^35, 36^. Lastly, differences in mouse genetic backgrounds appear to have contributed substantially to the quantitative evolution of the Ag-specific IgG concentrations with time.

### Validation of the sensor coating density in BioLayer Interferometry

The rising response of IgG with time during Year 1 in BALB/c mice suggests affinity maturation upon a single injection of a submicrogram dose of Ag. To better understand these long-lasting IgG responses quantitatively, we used BioLayer interferometry to directly measure the binding between serum IgG and its target Ag. In our experimental design, a biotinylated Ag was initially loaded onto a streptavidin sensor, and then the sensor was dipped into sera diluted in Buffer 1 (Materials and Methods) to monitor the binding signals in real time. IgG have two Fabs for Ag binding, with bivalent binding of the Ag leading to slower off-rate from the sensor. To assess affinity maturation, it is essential to coat the streptavidin sensor with biotinylated Ag at a low surface density to ensure measurement of monovalent binding and dissociation by IgG. To calibrate surface coating density, we first measured binding using a purified commercial monoclonal IgG2a specific for SARS-CoV-2 RBD (BioLegend #944803, mAb1 herein) at three different coating concentrations of biotinylated RBD (Biotin-RBD): 20, 50 and 100 nM. Moreover, the Biotin-RBD was prepared by site-specific biotinylation using the free engineered cysteine near the C-terminus of the RBD (Materials and Methods) to ensure that the epitope on RBD was unmodified. Fig. 4a-b show these binding and dissociation sensorgrams for mAb1. All the dissociation traces in Fig. 4b can be well described by single exponential functions (black lines), which yielded the rate of dissociation for IgG from sensors coated with different concentrations of Biotin-RBD. These observed rate constants from two independent repeats of the same experiments were then plotted in Fig. 4e (black circles) as a function of the concentration of biotin-RBD that was used to coat streptavidin sensors. For these rate constants, a one-way ANOVA test returned a p-value of 0.03428, demonstrating that at the 0.05 level, the population means are significantly different. Specifically, the rate of dissociation from the 100-nM biotin-RBD surface was lower than that of either 20 nM or 50 nM biotin-RBD surface. This result is consistent with bivalent binding by IgG on streptavidin sensors coated with 100 nM biotin-RBD, but monovalent binding on streptavidin sensors coated with either 20 nM or 50 nM biotin-RBD.

**Fig 4.**
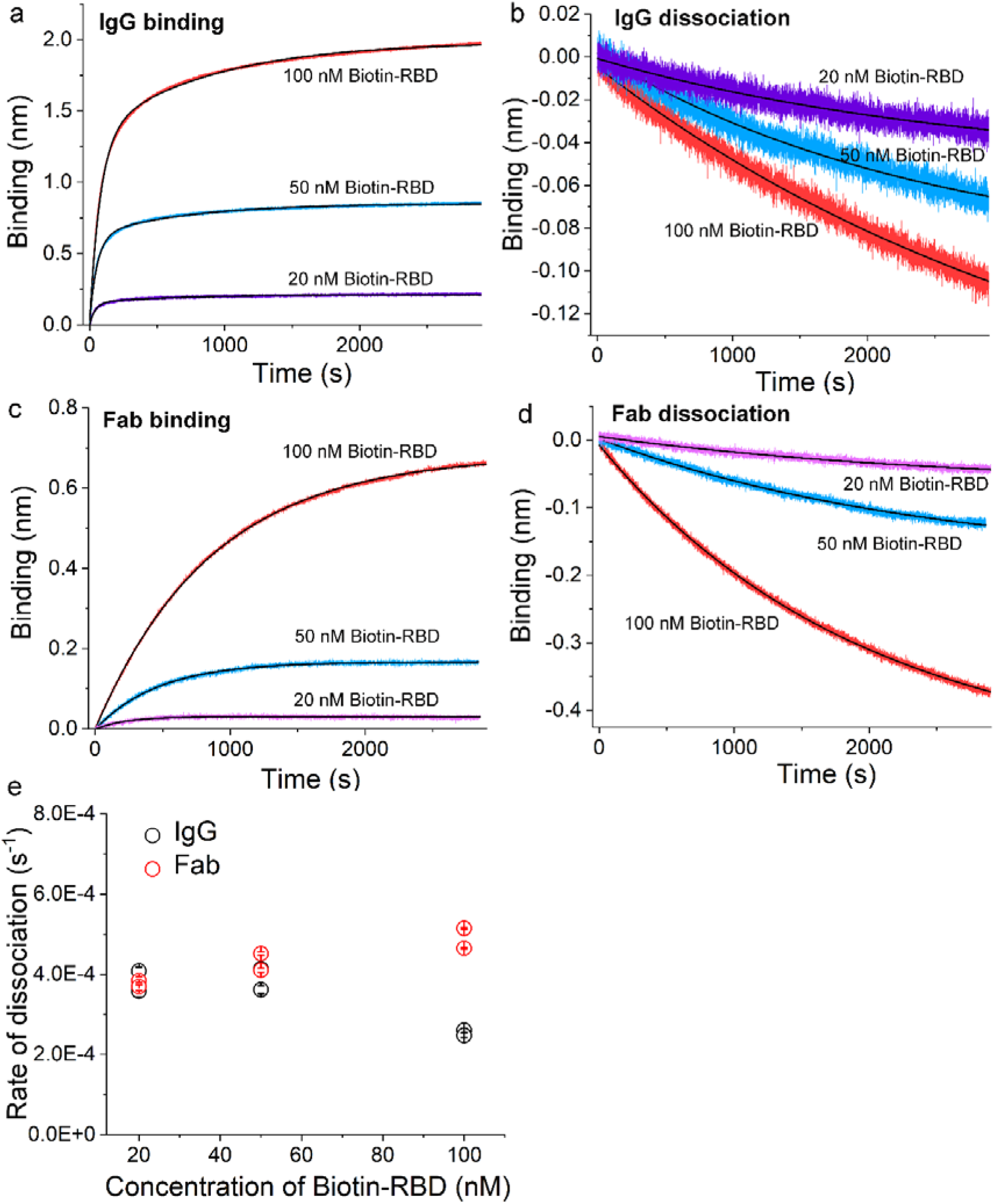
Validation of low-density coating of streptavidin sensors in BioLayer interferometry experiments. (**a**)(**b**) Real-time binding for mAb1 to biotin-RBD and its dissociation measured using BioLayer interferometry. The concentration of the IgG was 38 nM. (**c**)(**d**) Real-time binding for a Fab fragment to biotin-RBD and its dissociation measured using BioLayer interferometry. The concentration of the Fab was 62 nM. (a) through (d), the solid black lines were fits to both the binding and dissociation curves using exponential functions. (**e**) The observed rate constant of dissociation was plotted as a function of the concentration of biotin-RBD that was used to coat streptavidin sensors.

To confirm this, we prepared Fab fragments from mAb1 (Materials and Methods) and repeated the binding experiments above at the three different coating concentrations for streptavidin sensors, as shown in Fig. 4c-d. All the dissociation traces in Fig. 4d can be well described by single exponentials (black lines), which yielded the rate of dissociation for Fab fragments from sensors coated with different concentrations of Biotin-RBD. These observed rate constants from two independent repeats of the same experiments were then plotted in Fig. 4e (red circles) as a function of the concentration of biotin-RBD that was used to coat streptavidin sensors. Among these dissociation rate constants for Fab, a one-way ANOVA test returned a p-value of 0.05601, demonstrating that at the 0.05 level, the population means are not significantly different. Specifically, the rate of dissociation from the 100 nM biotin-RBD surface was statistically indistinguishable from that of either the 20 nM or the 50 nM biotin-RBD surface. This result is consistent with monovalent binding by Fab fragments on the streptavidin sensor throughout all three coating conditions. Furthermore, a one-way ANOVA test on the following five sets of rates of dissociation returned a p-value of 0.06043: IgG dissociation from sensors coated with 20 nM biotin-RBD, IgG dissociation from sensors coated with 50 nM biotin-RBD, Fab dissociation from sensors coated with 20 nM biotin-RBD, Fab dissociation from sensors coated with 50 nM biotin-RBD, and Fab dissociation from sensors coated with 100 nM biotin-RBD. This comparison shows that at the 0.05 level, none of these rate constants are significantly different from one another, and thus demonstrates that IgG binds to sensors coated with either 20 or 50 nM biotin-RBD in a monovalent manner. As a consequence, we have chosen 50 nM biotin-RBD or 50 nM biotin-HEL to coat streptavidin sensors for all BioLayer interferometry experiments.

### Ag-specific binding kinetics measured using BioLayer interferometry

With the sensor coating density established above, the sera from individual animals at different times post-immunization were then diluted in Buffer 1 and subjected to binding and dissociation measurements with their respective Ags in real time. Representative traces are shown in Fig. 5a and Fig. 5b for sera from a BALB/c mouse immunized with pRBD(DNA1)1 and in Fig. 5c and Fig. 5d for sera from a BALB/c mouse immunized with pHEL(DNA1)1, respectively. The amplitudes from real-time binding and dissociation allowed us to calculate the percentages of Ag-specific IgG that remained bound to Ags at the end of each dissociation period (Supporting Fig. S1a and S1b), which is a direct indicator of Ab affinity that is largely independent of their input concentrations (Supporting Fig. S1c). As shown in Fig. 5e, the percentages of Ag-specific IgG that remained bound at the end of dissociation showed a progressive increase until 2 months after the single injection with either pRBD(DNA1) (red symbols) or pHEL(DNA1) (blue symbols), demonstrating affinity maturation. These quantitative trends were in sharp contrast to that for the sera from a B6 gene knockout mouse that was deficient in both alpha/beta and gamma/delta T-cell receptors (TCR^-/-^)^37^ and immunized with pHEL(DNA1)1 (green diamonds), which served as a negative control for affinity maturation. Moreover, it is surprising to note that for all cases, regardless of the specific Ags, the off-rates of the Ag-specific IgG induced by either pRBD(DNA1) or pHEL(DNA1) were already at (5.2±0.3)×10^-4^ s^-1^ on Day 5 (D5) (red and blue symbols in Fig. 5f), including the TCR^-/-^ mouse (green diamonds in Fig. 5f). These values of off-rates were comparable to the off-rate of (3.9±0.3)×10^-4^ s^-1^ measured for mAb1, the commercial and affinity-matured mouse monoclonal IgG2a for RBD (Fig. 4e), which is shown as a dashed line in Fig. 5f. This commercial IgG2a can potently block the binding of RBD to human ACE2 at a concentration of 0.05-0.25 µg/mL (BioLegend CAT#944803). Over the 6-month period after a single injection, the off-rates of the Ag-specific IgG from all these animals underwent little or no changes (Fig. 5f). Thus, the affinity maturation for anti-RBD IgG as indicated in Fig. 5e may be driven primarily by optimization of the Ab on-rate^38, 39^.

**Fig. 5.**
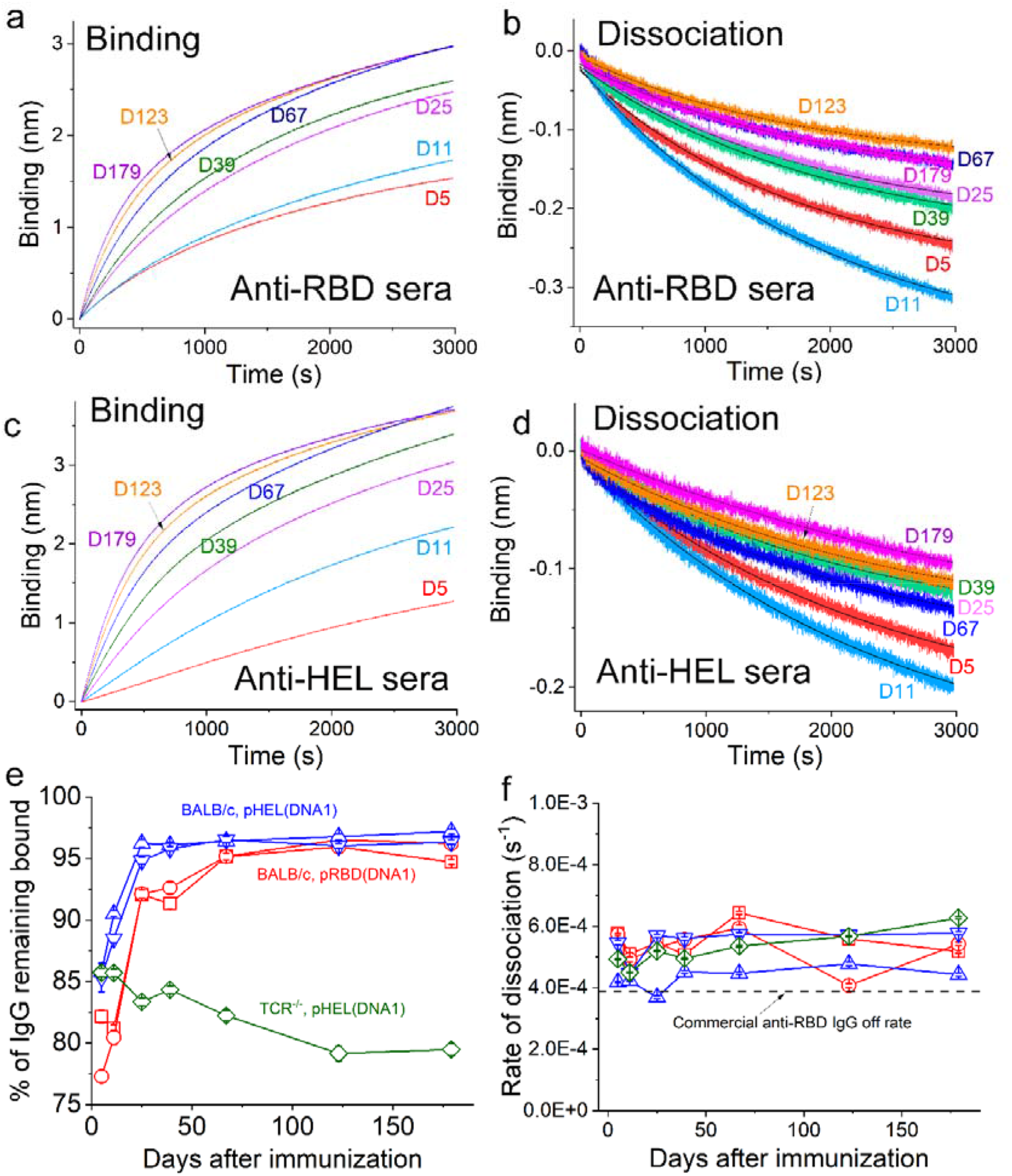
Kinetics for the binding and dissociation of various Ag-specific sera samples monitored using BioLayer interferometry. (**a**)(**b**)(**c**) and (**d**) Real-time binding and dissociation measurements for mouse sera from different days after immunization using SVLS as described in the text. (**e**)(**f**), percent of IgG remaining bound at the end of dissociation (e) and the dissociation rates (f) as a function of days after immunization measured using BioLayer interferometry for sera samples from various immunization conditions as indicated in (e). For both (e) and (f), red squares, red circles, upper triangles, down triangles and diamonds represent sera samples from immunizations using pRBD(DNA1)3, pRBD(DNA1)4, pHEL(DNA1)1, pHEL(DNA1)2, and pHEL(DNA1)1 respectively. The doses of RBD and HEL were 0.24 and 0.1 µg per animal, respectively. N=4 for each time point in (e) and (f).

### SVLS induce IgG affinity maturation in BALB/c mice

To confirm this, we sought to verify the concentration of Ag-specific IgG in mouse sera (Fig. 2) using an approach that is orthogonal to ELISA. The accurate concentration will then allow us to determine the on-rates without ambiguity. To this end, we first determined the concentration of Ag-specific IgG using a biotinylated Ag pulldown assay with streptavidin-coated magnetic beads (S-beads) followed by quantitative western blotting. These results are shown in Fig. 6a for the capture of RBD-specific IgG on D11 and D123, respectively, from a BALB/c mouse immunized with a single dose of pRBD(DNA1)1.

**Fig. 6.**
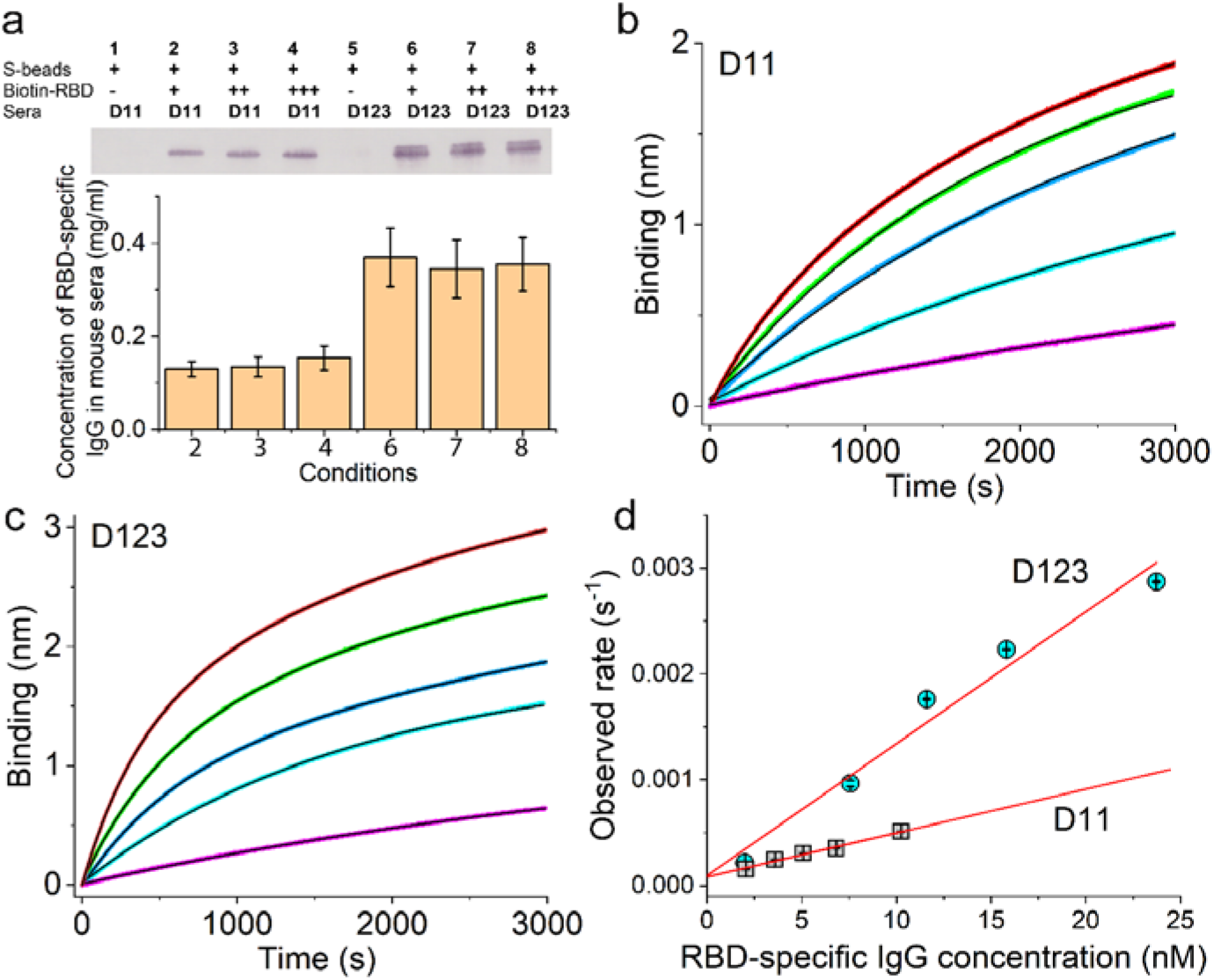
Affinity maturation of the RBD-specific IgG response induced by pRBD(DNA1) in BALB/c mice. (**a**) RBD-specific IgG capture using streptavidin-coated magnetic beads together with biotin-RBD. (**b**)(**c**) Real-time binding curves between biotin-RBD and mouse sera at various dilutions, with (b) for sera collected on D11 and (c) for sera collected on D123 after the single injection. The dose of RBD was 0.24 µg per animal. The concentrations of RBD-specific IgG in each dilution are shown in panel (d) along the x-axis. The exponential fits are shown in black solid curves. (**d**) The observed binding rate constants with standard errors obtained from exponential fits of the binding curves shown in (b) and (c) are plotted as a function of RBD-specific IgG concentration for sera from D11 (grey squares) and sera from D123 (cyan circles). Adjusted R-square values from linear regressions are 0.987 for D11 and 0.960 for D123, respectively.

To confirm complete capture of the RBD-specific IgG, we used increasing concentrations of biotin-RBD, as indicated on Lanes 2, 3 and 4 for D11 sera and Lanes 6, 7, and 8 for D123 sera (Fig. 6a top panel). Comparison with a standard curve (Supporting Fig. S2) obtained from the same western blots yielded the RBD-specific IgG concentrations for all these samples as shown in the bottom panel of Fig. 6a. For D11 sera, Lanes 2, 3 and 4 yielded very similar concentrations, with an average of 0.139±0.007 mg/mL. A one-way ANOVA test among Lanes 2, 3, and 4 returned a p-value of 0.717; therefore, there were no statistically significant differences among these measurements with increasing biotin-RBD, which indicates that we captured nearly all RBD-specific IgG from D11 sera. Also, this concentration was much higher than that measured by ELISA (Fig. 2b), suggesting that the conventional ELISA significantly underestimates the concentration of anti-RBD IgG on D11. For D123 sera, Lanes 6, 7 and 8 also yielded very similar concentrations, with an average of 0.356±0.007 mg/mL. A one-way ANOVA test among these lanes returned a p-value of 0.96, again demonstrating no statistically significant differences among these measurements with increasing biotin-RBD and indicating that we have captured nearly all RBD-specific IgG from D123 sera. However, a one-way ANOVA test between D11 and D123 measurements returned a p-value of 2.9e-5, indicating that the concentrations of RBD-specific IgG were significantly different in the sera from these two different dates.

These concentrations allowed us to measure the on-rates of RBD-specific IgG to RBD using serial dilutions of the respective sera in BioLayer interferometry, as shown in Fig. 6b for the series of binding curves from D11 sera and Fig. 6c for the series of binding curves from D123 sera (note the different scales of the y-axes in Fig. 6b versus 6c). All these binding curves can be well described by exponential functions (solid black lines). We plotted the observed rate constants from these exponential fits as a function of the RBD-specific IgG molar concentration in Fig. 6d, which showed the expected linear dependence for both D11 (grey squares) and D123 sera (cyan circles), with adjusted R-square values of 0.987 for D11 and 0.960 for D123, respectively (straight red lines). The slopes of these lines determined the bi-molecular rate constants for the binding between RBD and RBD-specific IgG, which were (4.2±0.2)×10^4^ M^-1^s^-1^ on D11 and (1.2±0.1)×10^5^ M^-1^s^-1^ on D123. Therefore, the on-rate increased almost 3-fold over the period from D11 to D123, demonstrating affinity maturation.

## Discussion

### (1) The minimal molecular determinants for a durable Ab response

Acute live viral infection in mice can lead to long-lasting antiviral serum Abs in both B6 and BALB/c mice^40^. At present, the mechanistic linkage between immediate and long-term Ab production upon viral infection or vaccination has not been fully established^41, 42^. This is due in part to the inherent complexity of studies of viral infections, as well as the empirical use of adjuvants or the complex ingredients of commonly studied VLPs. In many cases, a single injection of an immunogen in the absence of adjuvants is expected to produce a short-term Ab response only. In light of this background, it is notable that the IgG responses to a single immunization of SVLS at a very low Ag dose were long-lived, regardless of the displayed protein Ag or the mouse genetic background. In BALB/c mice, a single immunization with SVLS leads to lifelong secretion of IgG for both RBD and HEL. These results indicate that oriented protein display on a virus-sized structure, in the absence of any other adjuvants or Toll-like receptor (TLR) activation, can deliver long-term survival signals for Ag-specific plasma cells, true for both B6 and BALB/c mice. The presence of iNA can profoundly modulate the magnitude of the IgG response in both the short and long-term. These data thus established the minimal determinants of virus particles that give rise to long-lasting antiviral Ab responses from a primary response in mice. Overall, these results support the models in literature that Ag valency together with antigenic threshold may determine the durability of a protective Ab response^7^.

While the detailed molecular and cellular mechanisms that lead to the production of long-lived IgG response remain to be discovered, the differences in the long-term kinetic trends of IgG between the two mouse strains are intriguing. These data suggest that within the same animal species (mice), the durability of an IgG response towards the same immunogen at the same Ag dose can be modulated by differences in host genetic backgrounds, the detailed mechanisms of which remain to be unravelled. Along this line of strain differences, we noted recently that BALB/c mice mounted much stronger IgG1 than B6 on D11 for both pRBD and pHEL^19^, with all other immunization conditions being identical. These results suggest that early on in an Ab response to virus-like immunogens (between D5 and D11), BALB/c mice amplify the activation signals conveyed by epitope density more than B6. Whether this difference observed on D11 may lead to differences in IgG long-term trends between the two strains remains to be studied. Moreover, what causes these differences between B6 and BALB/c towards the same SVLS? Among several possibilities, whether these differences are imparted from the original Ag exposure^7^ (differences in B cell precursor frequency for example) or due to differences that occur at a later stage after the initial B cell activation and MHC class II Ag presentation^43^ remain to be tested.

### (2) Virus-like immunogens elicit Ag-specific IgG with exceptional off-rate

Another surprising result from the current study is that these Ag-specific IgG elicited by SVLS exhibit very slow off-rates that are less than 0.001 s^-1^, which are comparable to that of a commercial affinity-matured mAb (Fig. 5f). These IgGs emerged as early as Day 5 post Ag exposure, regardless of the specific protein Ag or the mice strains and even in the absence of functional T cells (Fig. 5f). The IgG from both B6 and BALB/c mice can potently neutralize cognate pseudovirions (Fig. 2c). These data are consistent with previous studies of live infection by vesicular stomatitis virus (VSV) and influenza virus in murine models. It has been found that VSV-specific IgG in germline configuration is already highly protective against VSV infection^44^, which does not require somatic hypermutation or affinity maturation^44^. Similar results also occur in murine models of influenza virus infection^45^. The current study offers a mechanistic explanation for these live infection data: (1) live infection produces virions; (2) the consensus features of virion structures, i.e., epitope density (ED) and iNA in the context of a virion-sized structure, can robustly activate naïve Ag-specific B cells to initiate class-switch recombination and secrete IgG that is independent of viral replication or any other adjuvants; and (3) the IgG has an intrinsically slow off-rate that can offer effective neutralization of cognate virions. The slow IgG off-rates that we observed even for a nonviral protein, HEL (Fig. 5f) suggests that this may be a phenomenon associated with IgG induction by virus-like immunogens in mice.

An optimal, highly potent IgG should have both a fast on-rate and a very slow off-rate so that it can bind its target with high affinity. Despite a very slow off-rate, this early version of RBD-specific IgG is not optimal because of a slow on-rate. As shown in Fig. 6d, the RBD-specific IgG on D11 post immunization has an on-rate of (4.2±0.2)×10^4^ M^-1^s^-1^, which is more than tenfold slower than the on-rate of the commercial RBD-specific mAb1 that we have measured, (6.1±0.3)×10^5^ M^-1^s^-1^ (data not shown). However, we hypothesize that this immediate IgG response is at evolutionary advantage to the host because defects in IgG on-rate can be potentially compensated for by a high concentration of the IgG rapidly secreted in the context of an extra-follicular response to acute viral infection, which is triggered by the combined signals from ED and iNA presented by intact virion structures^19^. This suggests that upon germinal center formation in a suitable host, the major improvements conferred by affinity maturation to these IgGs are likely in the on-rates of Ag binding. In fact, this is the case in our data (Fig. 6d): over a period of 2 months, the on-rate of the RBD-specific IgG increased by roughly 3-fold from the single injection. Also, in murine models of live infection by VSV, it was discovered that the efficiency of virion neutralization is highly correlated with the on-rates of viral-specific IgGs identified throughout the live viral infection^39, 46^, suggesting that this optimization of IgG on-rate is indeed the case for live viral infections. In a separate study, enhancement of IgG on-rate during affinity maturation was also observed for human nAbs induced upon infection by the respiratory syncytial virus^38^.

Therefore, in both mice and humans, live infections of two different viruses optimized the on-rate of IgG, suggesting that the germline progenitor of this IgG has excellent off-rates to provide effective protection. Taken together, this is a highly effective strategy for antiviral defense. The speed and magnitude of this IgG response are largely independent of T cells, affording the host with a mechanism for a fast and strong nAb response that would be critical during the early onset^47^ of an antiviral defense. It’s important to note that, in addition to binding rates, an IgG’s potency for neutralizing viral infectivity is also influenced by the virus’ ED. Specifically, the ED of the virus can impact the stability and effectiveness of bivalent binding by the IgG, which is more effective than monovalent binding^48, 49^.

### (3) SVLS as new platform for dissection of antiviral response and potential vaccines

In comparison to the complex bacteriophage VLPs that have been used in studies of Ab responses to virus-like immunogens, SVLS use the minimal essential viral components to model these Ags, yet the potency of the IgG response triggered by these SVLS can be compared to that of well-established bacteriophage VLPs^19^, suggesting that this platform has captured the key features of VLPs which trigger an Ab response. In current study, we show that a single immunization with SVLS at low Ag doses in mice can lead to long-lasting nAb responses, which is highly desired for antiviral vaccines. It remains to be tested if this phenomenon of *‘single injection with lifelong Abs’* as we observed in BALB/c mice is applicable to higher order mammalian species. If so, this system of SVLS will be of tremendous value in our understanding of human Ab response and also hold great promise for the development of various clinically relevant vaccines without the use of additional adjuvants. This platform is highly modular and can be readily adapted to different proteins.

We have tested RBD and HEL as shown in the current study and a self-Ag peptide previously^23^. Therefore, this platform has the potential for use in other viral protein systems for understanding Ab responses without an active infection, as well as potential applications in vaccines. The modular nature of the SVLS structure is particularly desirable for dissecting the roles of individual components in Ab responses towards complex immunogens such as virions. The completely synthetic nature of these SVLS would be useful for vaccine design and formulation from several different perspectives. First, the ability to use iNA for intrinsic TLR activation can dramatically enhance B cell activation^19^, which may be strongly desired for activating those very rare Ag-specific B cells. In this regard, SVLS may be at an advantage compared to lumazine synthase^50^, ferritin nanoparticles^51^ or computationally designed self-assembling protein nanomaterials^52^. Second, SVLS use a stable formulation^17,18^ to preserve the integrity of virus-like structures for presentation of protein Ags in their native conformations without any associated viral infectivity, which may be at an advantage compared to inactivated viruses, where formaldehyde and/or detergent treatment can alter conformations of protein Ags or even destroy the structures of viruses that are critical for B cell activation^7^. Third, the regulatory requirements for purified synthetic agents without exogenous adjuvants could be less complex compared to products directly derived from cell culture^53^, and last, the use of nucleic acids with native phosphodiester backbones in SVLS without the need for nuclease-resistant phosphorothioate could substantially reduce the cost of vaccine manufacture.

## Conclusion

Using mouse as our current model system, we have identified the minimal components and features present in typical enveloped virions that can give rise to long-lasting antiviral IgG in a primary immune response in the absence of exogenous adjuvants. This study helps understand the durability of antiviral antibody responses and has implications on vaccine design to generate durable antibody responses.

## MATERIALS AND METHODS

### Synthesis of SVLS

All SVLS used throughout this study were prepared following protocols we published previously^17, 18, 23^. Specifically, three different lipids of designated molar ratios were used in the synthesis of liposomes as we described^17^: 1,2-distearoyl-sn-glycero-3-phosphocholine (DSPC) (Avanti Polar Lipids, >99% purity), 1,2-distearoyl-sn-glycero-3-phosphoethanolamine-N-[maleimide (polyethylene glycol)-2000] ammonium salt (DSPE-PEG (2000) maleimide) (Avanti Polar Lipids, >99% purity) and cholesterol (Avanti Polar Lipids, >98% purity). The choice of a neutral lipid DSPC is meant to mimic that of cell membranes, from which enveloped viruses derive their viral membranes^26^. For all SVLS in current studies, DSPC and cholesterol comprised ≥ 99% of all lipids. These structures display protein antigens (Ags) of choice in a specific orientation on their surfaces with a regulated ED that quantitatively mimics the surface of naturally occurring viruses^48, 54^. The SARS-CoV-2 RBD recombinant protein used in this study was overexpressed in house using 293F cells and purified to >95% purity following our established protocols^18^. The HEL recombinant protein used in this study contains two site-directed mutations at R73E and D101R and was overexpressed in *E. coli* and purified to >95% purity in house following our established protocols^17^. DNA1 was custom synthesized by IDT. The sequence of DNA1 is as follows, 5’-TCCATGA<u>CG</u>TTCCTGA<u>CG</u>TT-3’. The calculated molecular weight of DNA1 is 6059.0 Dalton. The measured molecular weight of DNA1 by electrospray ionization mass spectrometry is 6058.5 Dalton. Epitope density (ED) was quantified using methods that we established previously ^17, 18, 23, 55^ that were also validated by single-molecule fluorescence techniques that were developed and established in house^56, 57^. For SVLS with iNA, the average number of iNA molecules per SVLS was also quantified using methods that we established previously^17, 18^.

### Mice immunizations

All animal procedures were approved by the University of Michigan Animal Care and Use Committee. Female C57BL/6 or BALBc/J mice (8 weeks, Jackson Laboratory) were used for immunizations. Prior to inoculation, all injection samples were filtered through a 0.45 µm pore size membrane to eliminate potential microbial contamination. 100 µL samples at a dose between 0.1 and 0.48 µg of respective protein Ags were injected into each mouse subcutaneously, 50 µL in each flank. Throughout the entire study, no adjuvants other than the SVLS were administered unless otherwise noted and only a single injection was administered throughout. Mouse blood was collected submentally using Microvette serum collection tubes (Sarstedt) three days before the first injection, and 5 days thereafter following specified blood collection schedules. The sera were harvested by centrifugation at 10,000 g for 5 minutes, aliquoted and immediately frozen and stored at -80°C. The gene knockout mice Tcrb^tm1Mom^/ Tcrd^tm1Mom^ (#002122) (TCR^-/-^ herein) in B6 background were deficient in both alpha beta and gamma delta T-cell receptors^37^ and were purchased from Jackson Laboratory. Gene knockout mice were housed in a germ-free environment and immunization protocols were carried out as described above.

### Enzyme-Linked Immunosorbent Assay (ELISA)

Blood serum was tested by ELISA in order to quantitate Ag-specific IgG responses to various immunizations. 96-well plates (Nunc MaxiSorp, Invitrogen) were coated overnight at 4°C with either 200 ng of sHEL or 320 ng of sRBD per well in PBS. After blocking with 1% Bovine Serum Albumin (BSA, Fisher) in PBS, mouse sera of specified dilution factors were added to each well for incubation at 22°C for 2 hours. After three washes using PBS with 0.05% Tween 20, secondary goat anti-mouse IgG Fc-HRP antibody (# 1033-05, SouthernBiotech) was added in blocking buffer at 1:6000 dilution and incubated for 1 hour at 22°C. Following three washes, 100 µL of substrate 3,3’,5,5’-Tetramethylbenzidine (Thermo Scientific) was added to each well and incubated in the dark for 10 minutes. The reaction was stopped by addition of 100 µL 2M sulfuric acid in each well. The optical density of each well at 450 nm was measured using a microplate reader (Bio-Tek Synergy HT). All the OD values reported were background subtracted by comparison between two wells that were coated with soluble protein and PBS, respectively. To estimate Ag-specific IgG concentrations in mouse sera based on ELISA standard curves, all sera samples were properly diluted to reach final OD values between 0.2 and 0.5 before interpolation. Specifically, we used mAb1, the commercial RBD-specific monoclonal mouse IgG2a (BioLegend CAT#944803) as the known reference to construct standard curves for detection of RBD-specific IgG and IgG2a. We used mAb2, the HEL-specific monoclonal mouse IgG1 (clone HyHEL10, a special gift from Prof. Irina Grigorova) as the known reference to construct standard curves for detection of HEL-specific IgG and IgG1.

### Preparation of HIV-1 virions pseudotyped with SARS-CoV-2 envelope

The HIV-1 pseudotyped with SARS-CoV-2 envelope was prepared following published protocols ^58^ but with modifications. Briefly, HEK 293T/17 cells (ATCC, Manassas, VA) were cultured at 37°C with 5% CO_2_ in DMEM supplemented with 10% FBS (HyClone Laboratories, Logan, UT). 10^6^ 293T cells in a 2-mL culture volume were seeded overnight in a 35-mm dish before transfection using the TransIT LT-1 transfection reagent (Mirus Bio, Madison, WI). For each dish, 2 μg of the provirus-containing plasmid pNL4-3 luc R^-^E^-59^ (ARP-3418, NIH AIDS Research and Reference Reagent Program) was used to make the transfection reagent mixture, together with 1 μg of envelope expression plasmid pcDNA3.1 SARS-CoV-2 S D614G ^58^. The plasmid pcDNA3.1 SARS-CoV-2 S D614G was a gift from Jeremy Luban (Addgene plasmid # 158075; http://n2t.net/addgene:158075; RRID:Addgene_158075). The RBD amino acid sequence 328-537 encoded in this plasmid is identical to the sequence of RBD that we studied here. The transfection reagent mixture was incubated at room temperature for 15 min before drop wise addition to the culture medium, as we did previously ^60^. At 6 hours post transfection, the culture medium together with the transfection reagents was replaced with fresh complete medium and the incubation was continued at 37°C with 5% CO_2_. At 48 hours post transfection, the entire culture medium containing single-cycle HIV-1 viruses was collected and filtered through a 0.45-µm syringe filter (Millex-HV PVDF, Millipore). The filtrate was then aliquoted on ice, flash-frozen in liquid nitrogen and stored in a -80°C freezer. The concentration of virion particles was quantitated using a HIV-1 p24 ELISA kit (CAT#XB-1000, XpressBio) as we described previously ^60^.

### Virion neutralization assay

The virion neutralization assay follows the protocols we established previously but with important modifications ^60^. HIV-1 virions pseudotyped with SARS-CoV-2 envelope containing 9 ng of HIV-1 p24 were incubated with various mouse sera at 37°C for one hour, and then diluted with complete medium by 8-fold in volume for the sera to initiate infection of Huh-7.5 cells at 37°C for 2 hours. At the end of 2 hours, fresh medium was added to each well in a 24-well plate and the incubation was continued at 37°C with 5% CO_2_. Luciferase activity was measured 48 hours after infection. Briefly, the culture medium was removed and replaced with 100 μL of complete medium. 100 μL of Bright-Glo^TM^ reagent (CAT#E2610, Promega) that was just warmed to room temperature was then added to each well. The cells were incubated for three minutes at room temperature to allow cell lysis. After three minutes, 100 μL of lysate from each well was transferred to a single well in a 96-well black microtiter plate (Costar). Luminescence was measured using a Synergy^TM^ HT multi-mode plate reader (BioTek Instruments Inc., Vermont) and background luminescence was subtracted using Huh-7.5 cells without virus infection. For comparison among different time points post immunization, the luminescence readings for cells incubated with sera from each group of mice before injections was set as 100%. The luminescence readings from other time points for the same group were all normalized based on this and plotted as a percentage of remaining infectivity.

### BioLayer interferometry measurements

Affinity maturation of Ag-specific antibodies in animal sera together with binding and dissociation rate constants were measured using BioLayer interferometry. Briefly, the streptavidin sensor (Sartorius Corp) was coated with 50 nM Biotin-RBD or Biotin-HEL in 1×PBS for 30 min at 20°C, which was followed by washing in PBS for 10 min. For this application, both RBD and HEL were site-specifically biotinylated in house using EZ-link maleimide-PEG2-biotin (ThermoFisher, CAT#A39261) so that the binding epitopes on these proteins were not modified. After sensor loading, they were dipped into antibodies or sera of varied dilutions in Buffer 1 to measure the binding in real time using an OctetRed BioLayer Interferometer equipped with 8 sensor positions that were read simultaneously. The measurement for binding was continued for the indicated time, which was followed by dipping the sensors into Buffer 1 for the indicated time to monitor the dissociation in real time. Throughout, the measurements were done at 20°C. A PBS coated sensor control was included and subtracted from all the kinetic data before quantitative analysis. Buffer 1 was 1×PBS with 0.05% Tween 20 and 0.1% BSA. The percentage of IgG remaining bound, p, was determined by using the amplitude information from both binding and dissociation curves, specifically as follows: p=(A1+A2)/A1*100%, in which A1, a positive number, is the final amplitude at the end of binding measurement, and A2, a negative number as the result of dissociation, is the final amplitude at the end of dissociation measurement, as illustrated in Supporting Fig. S1a and S1b. In the current study, most binding and dissociation events were measured continuously in real time for 50 min each. For experiments shown in Fig. 6, the binding and dissociation were monitored continuously for 4000 s for better definition of the rate constants. The control Fab was prepared from the IgG form of the antibody (mAb1) using a Pierce^TM^ Fab Micro preparation kit (ThermoFisher Scientific, CAT#44685) following manufacturer instructions.

### Measurement of Ag-specific IgG concentration in animal sera using magnetic beads

We measured the concentration of Ag-specific IgG in the sera of immunized animals using streptavidin-coated magnetic beads. Specifically, high capacity Magne streptavidin beads (Promega CAT#V7820) were first exchanged into PBS buffer, and then mixed with biotinylated RBD or HEL for binding at 20°C for 1 hr with constant mixing on a lab vortexer. After this, the beads were washed 3 times using Buffer 1 and then mixed with sera also diluted in Buffer 1. The binding was continued at 20°C for 2 hrs with constant mixing. Finally, the supernatant was removed, and the bound IgG was resuspended in nonreducing Laemmli sample buffer at 95°C for 10 min. The amount of Ag-specific IgG was then quantitated by running a western blot using mouse IgG (BioLegend CAT#944803) of known mass as a standard. The secondary antibody used was goat anti-mouse IgG (Fc specific) conjugated with alkaline phosphatase from Sigma (CAT#A1418) and used at 1:3000 dilution.

### Statistical analysis

Statistical analysis was carried out using the Statistics toolbox in MATLAB (Mathworks, MA). A comparison of two independent variables was carried out using a two-sample T-test. Data sets with more than two independent variables were analyzed using a one-way analysis of variance as indicated. P-values less than 0.05 were considered statistically significant. Throughout, all errors in the main text were reported as standard errors unless otherwise noted; all data points in figures were reported as mean ± standard error unless otherwise noted.

## ACKNOWLEDGEMENTS

This work was supported by an NIH/NIAID grant (1R01AI155653-01A1) to WC and JZ. ARM was supported in part by an NIH/NIGMS T32 GM145304 Cellular Biotechnology Training Program. We thank Prof. Irina Grigorova for critical reading of the work during the early development of the manuscript. We thank Prof. Marilia Cascalho for very helpful discussion regarding the data and this work. We thank Dr. Charlie Rice at Rockefeller University for the kind gift of Huh-7.5 cells. The following reagent was obtained through the AIDS Research and Reference Reagent Program, Division of AIDS, National Institute of Allergy and Infectious Diseases (NIAID), National Institutes of Health (NIH): pNL4-3.Luc.R-E-from Dr. Nathaniel Landau. The plasmid pcDNA3.1 SARS-CoV-2 S D614G was a gift from Dr. Jeremy Luban through Addgene.

## ABBREVIATIONS

SVLS: synthetic virus-like structures
Ab: antibody
Ag: antigen
NA: nucleic acid
iNA: internal nucleic acid
ED: epitope density
SARS-CoV-2: the severe acute respiratory syndrome coronavirus 2
RBD: receptor binding domain
HEL: hen egg lysozyme
pRBD: SVLS that display RBD on liposomal surface without iNA
pRBD(DNA1): SVLS that display RBD on liposomal surface and encapsulate DNA1 as iNA
pHEL: SVLS that display HEL on liposomal surface without iNA
pHEL(DNA1): SVLS that display HEL on liposomal surface and encapsulate DNA1 as iNA
DSPC: 1,2-distearoyl-sn-glycero-3-phosphocholine
DSPE-PEG maleimide: 1,2-distearoyl-sn-glycero-3-phosphoethanolamine-N-[maleimide (polyethylene glycol)-2000] ammonium salt.

## Table of Content Graphic

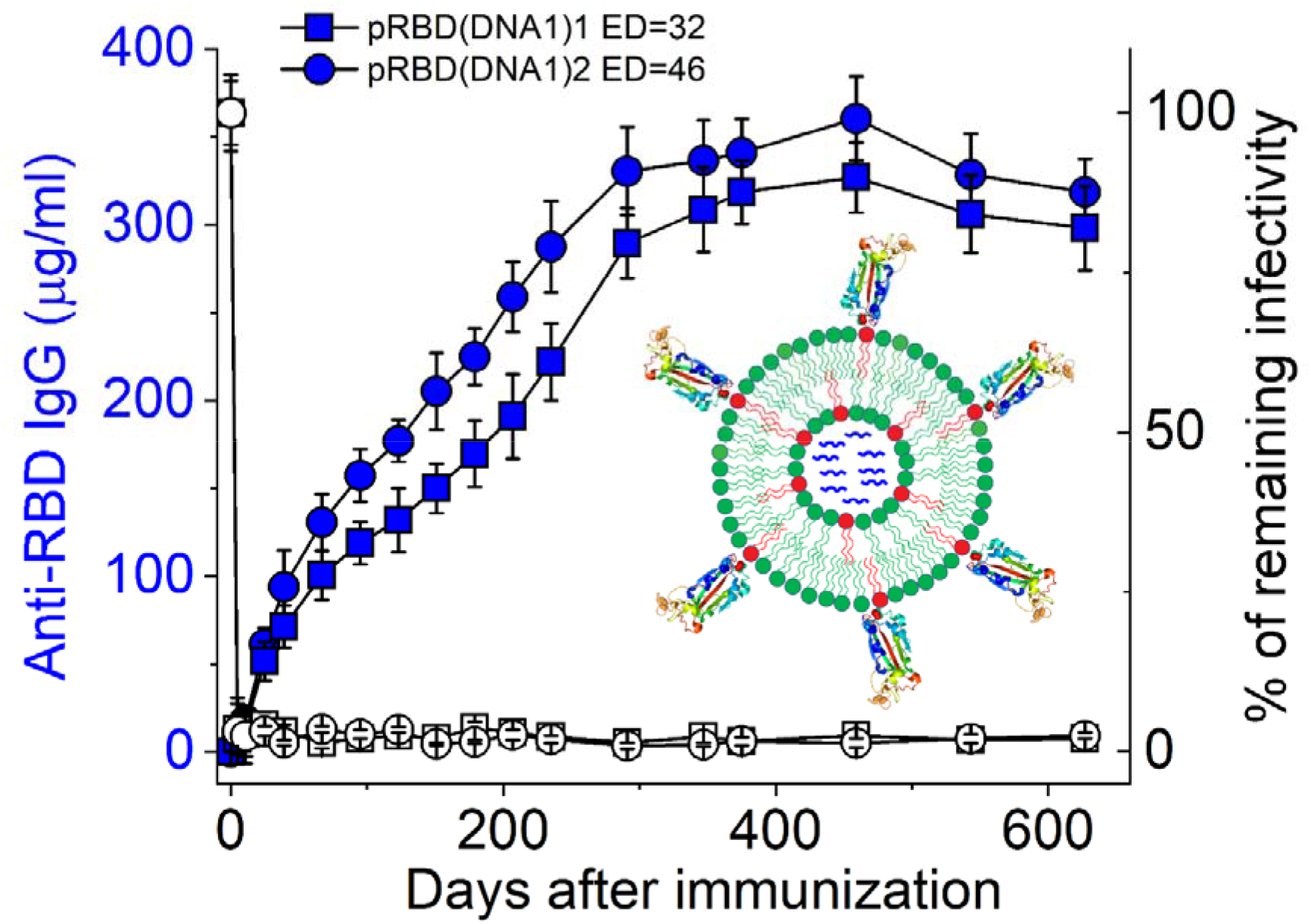

## Supporting Information

**Fig. S1.**
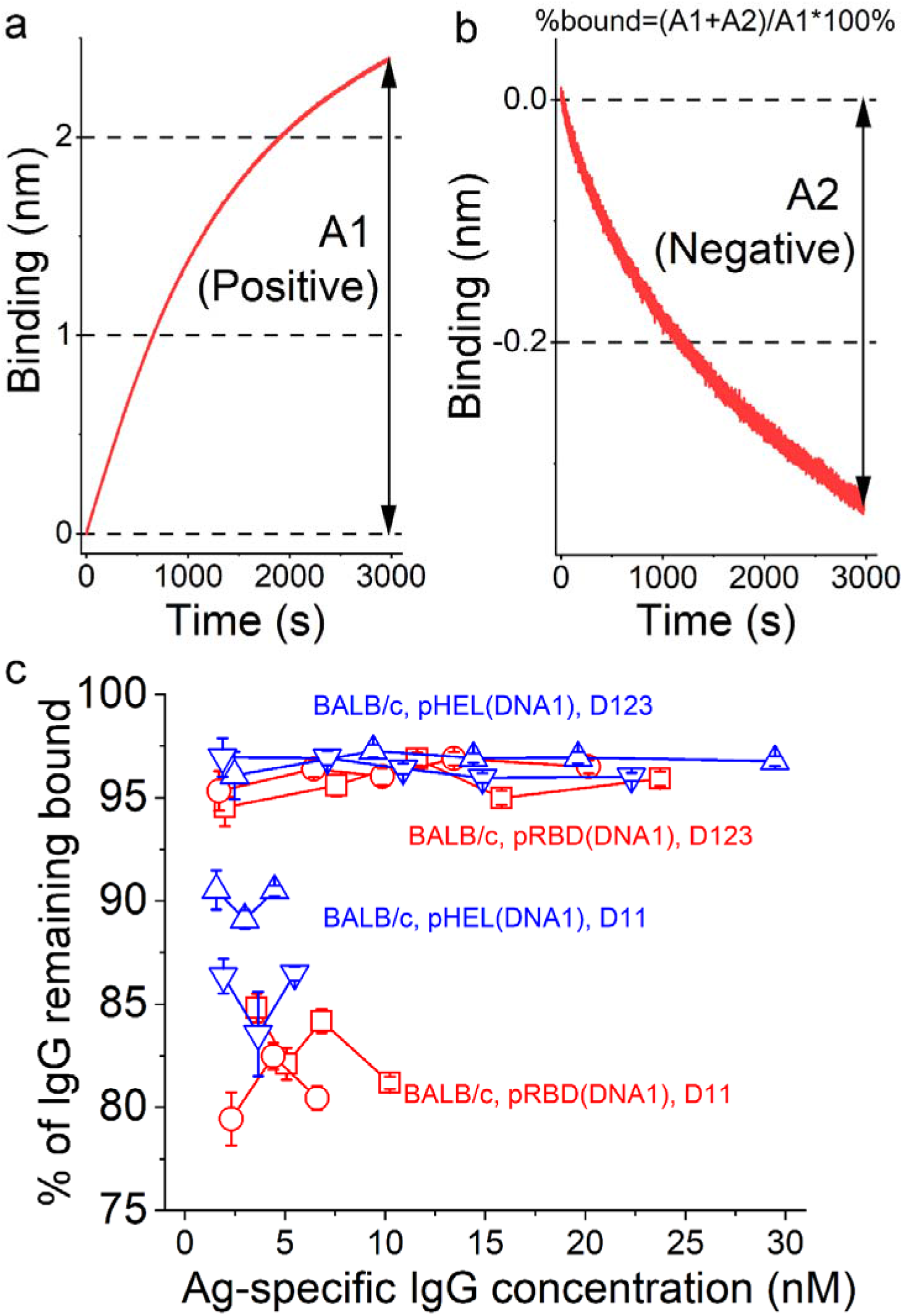
Indicator of affinity maturation that is largely independent of sera IgG concentrations. (**a**) A representative serum IgG binding curve showing the definition of binding amplitude, A1, that is measured from BioLayer interferometry experiments. A1 is the difference in the binding signal between the end and the beginning of a binding curve, which takes a positive value with a unit of nm. (**b**) The corresponding dissociation curve from (a) showing the definition of dissociation amplitude, A2, that is measured from BioLayer interferometry experiments. A2 is the difference in the binding signal between the end and the beginning of a dissociation curve, which takes a negative value with a unit of nm. The percentage of IgG remaining bound at the end of dissociation is calculated as indicated on top of panel (b). (**c**) The percentage of IgG that remains bound at the end of dissociation was plotted as a function of the Ag-specific IgG concentration. For each color and symbol, the different concentrations were realized by dilution of the sera and then BioLayer interferometry experiments were conducted. This was done for several serum samples from BALB/c mice immunized with the agents as noted in the figure. Specifically, red squares, red circles, upper triangles, and down triangles are for pRBD(DNA1)3, pRBD(DNA1)4, pHEL(DNA1)1, and pHEL(DNA1)2, respectively. The dose of RBD was 0.24 µg per animal and the dose of HEL was 0.1 µg per animal. The concentrations for Ag-specific IgG were determined using magnetic beads capture as described in Materials and Methods. N=4 for each Ag-specific IgG concentration in (c).

**Fig. S2.**
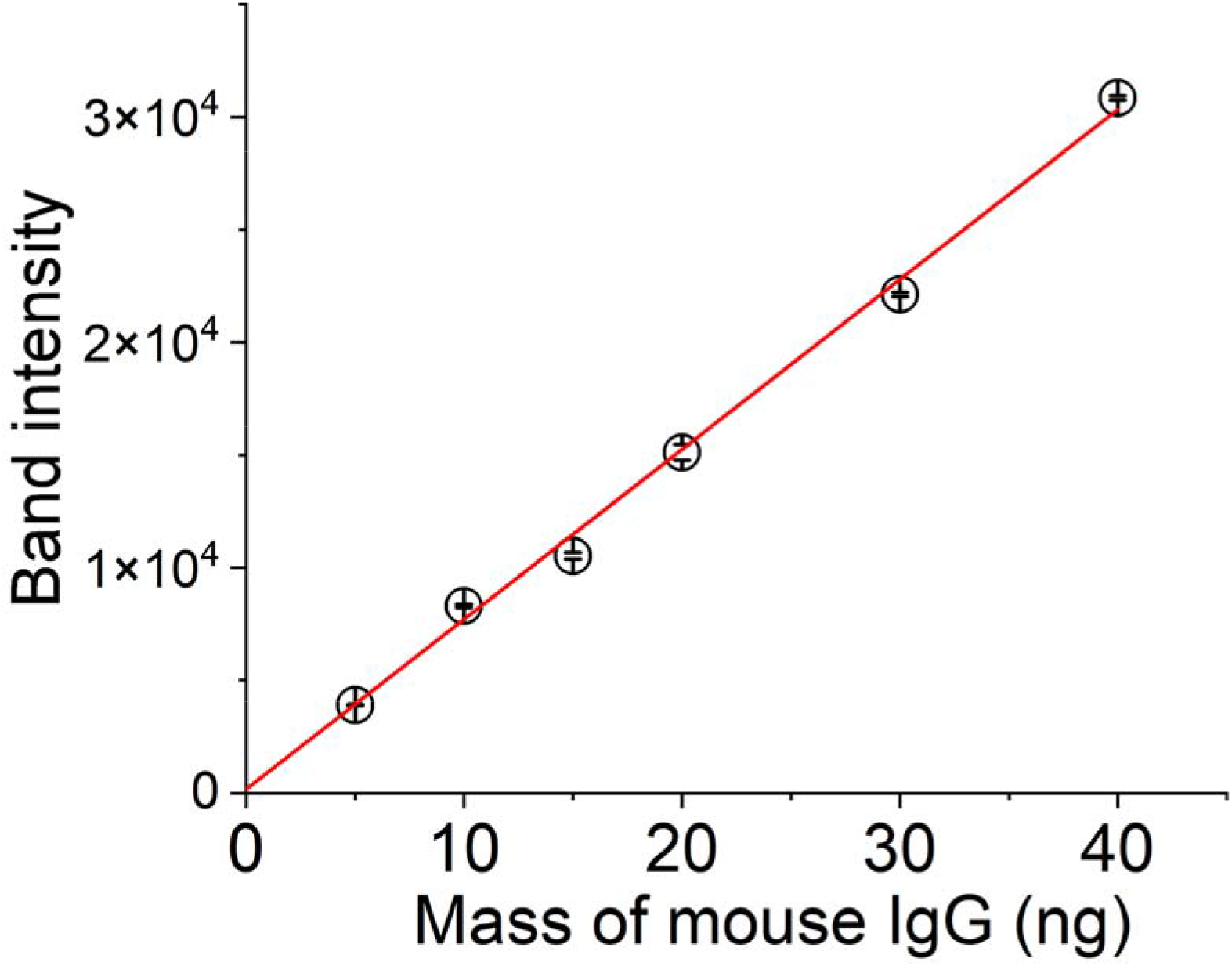
A representative standard curve from quantitative western blots to determine the concentration of Ag-specific IgG in mouse sera. We used mAb1, a commercial monoclonal mouse IgG (BioLegend CAT#944803) as the known reference to construct a standard curve on the same western blot on which Ag-specific IgG antibodies captured by streptavidin magnetic beads were loaded. The band intensity was quantified using ImageJ and plotted against the loaded references (N=3 for each data point). A linear regression (the straight red line) was then used to quantitate the mass of Ag-specific IgG in serum samples. The adjusted R-square value from the linear regression as shown above is 0.998. The mass was then used to calculate the concentration of the IgG based on volume dilutions.

